# Age-Related RPE changes in Wildtype C57BL/6J Mice between 2 and 32 Months

**DOI:** 10.1101/2024.01.30.574142

**Authors:** Debresha A. Shelton, Isabelle Gefke, Vivian Summers, Yong-Kyu Kim, Hanyi Yu, Yana Getz, Salma Ferdous, Kevin Donaldson, Kristie Liao, Jack T. Papania, Micah A. Chrenek, Jeffrey H. Boatright, John M. Nickerson

**Author notes:** Correspondence to: Please address all correspondence to Dr. John M. Nickerson; Department of Ophthalmology, Emory University, B5602, 1365B Clifton Road, NE, Atlanta, GA, 30322.

## Abstract

**Purpose:** This study provides a systematic evaluation of age-related changes in RPE cell structure and function using a morphometric approach. We aim to better capture nuanced predictive changes in cell heterogeneity that reflect loss of RPE integrity during normal aging. Using C57BL6/J mice ranging from P60-P730, we sought to evaluate how regional changes in RPE shape reflect incremental losses in RPE cell function with advancing age. We hypothesize that tracking global morphological changes in RPE is predictive of functional defects over time.

**Methods:** We tested three groups of C57BL/6J mice (young: P60-180; Middle-aged: P365-729; aged: 730+) for function and structural defects using electroretinograms, immunofluorescence, and phagocytosis assays.

*Results:* The largest changes in RPE morphology were evident between the young and aged groups, while the middle-aged group exhibited smaller but notable region-specific differences. We observed a 1.9-fold increase in cytoplasmic alpha-catenin expression specifically in the central-medial region of the eye between the young and aged group. There was an 8-fold increase in subretinal, IBA-1-positive immune cell recruitment and a significant decrease in visual function in aged mice compared to young mice. Functional defects in the RPE corroborated by changes in RPE phagocytotic capacity.

*Conclusions:* The marked increase of cytoplasmic alpha-catenin expression and subretinal immune cell deposition, and decreased visual output coincide with regional changes in RPE cell morphometrics when stratified by age. These cumulative changes in the RPE morphology showed predictive regional patterns of stress associated with loss of RPE integrity.

## 1. Introduction

The retinal pigment epithelium (RPE) is a monolayer of hexagonal cells that are located between the neurosensory retina and the choroid, that react with Bruch’s membrane ((1–5)). One of the RPE’s main functions includes supplying the retina with nutrients via basal infoldings and removing waste by-products from the photosensory processes that take place in the photoreceptor cells ((4,5)), specifically by recycling the tips of the outer segments of rods and cones with its apical microvilli via phagocytosis ((6)(7,8)). The RPE is also critical for maintaining homeostasis by secreting growth hormones to aid in maintaining the structural integrity of the choriocapillaris endothelium and the photoreceptor cells ((6)). The role of the RPE is evident beginning at the very early stages of ocular development. Ablation of the RPE before E10 in mice results in arrest of eye growth and resorption, while late ablation after RPE differentiation results in disorganization of retinal layers and ectopic routing of retinal ganglion cells ((9,10) The RPE also aids in immune regulation by secreting immunosuppressive factors ((6)). Due to the plethora of functions that the RPE performs, the density, structure, and function of the RPE is critical to maintaining homeostasis of the eye. Because each RPE cell can support up to 20-45 photoreceptors, even small changes in RPE structure or metabolomics can have effects on the function and development within the ocular environment [(10–15)(16,17). [(18,19)]. Thus, studying changes in RPE structure during aging can be a predictive metric by which to understand retinal degeneration and loss of vision. Due to the quiescent, post mitotic status of the RPE, its sensitivity to the high metabolic demands of photoreceptors, and its reduced capacity for regeneration after damage it is susceptible to degeneration and dysfunction that can affect the entire eye.

Aging affects many parts of the human body including the RPE. Even in healthy adults, RPE cells uniformly decrease in density from the second to the ninth decade of life ((20)), decreasing by about 0.3% per year with increasing age ((21)), which leads to a higher ratio of photoreceptors per RPE cell ((22)). Del Priore *et al.* discovered that the number of apoptotic RPE cells significantly increases with age in humans and were predominantly found at the macula which contains a higher density of photoreceptors than the periphery ((23,24,25)). A combination of increased photoreceptor-RPE ratio, failure of RPE cytokinesis, defects in RPE mitochondrial metabolic activity, and decreased damage repair that accumulates with age can impede the cells’ optimal bioenergetics, (26,27)ultimately contributing to loss of RPE function, (26,27).

Many other changes occur during aging of the RPE including the retinal pigment epithelium-Bruch’s membrane complex thickens at the foveal minimum, or center-point foveal thickness, an accumulation of lipofuscin, increased circulation of profibrotic macrophages, choroidal neovascularization, and the formation of hard drusen ((28–31) ((32,33). While each of these may occur in natural aging, there is also overlap in these characteristics during disease onset, suggesting at least a partially conserved mechanism is employed during normal and abnormal aging paradigms(34,35). (36)(6)Age is the strongest demographic risk factor for AMD, followed by other factors including race, iris color, dysregulation of genes associated with the immune system, and lipoproteins ((37,38). A wealth of data about AMD and genetic links to clinical phenotypes that characterize the disease have been gleaned from the AREDS (Age-related Eye Disease Study) and AREDS2. The goal of these randomized, clinical studies was to assess the effectiveness of high doses of antioxidants like beta-carotene to stabilize and enrich components of the visual cycle (which can help support aged RPE cells), low doses of zinc, and omega-3 fatty acids supplementation reduces the risk of AMD onset in patients over a 5-10-year period. Genome-wide association studies have linked an increased risk of AMD onset and poor visual acuity with single nucleotide polymorphisms in genes associated with RPE function and inflammation, like *CFH* and *ARMS/HRTA1,* and *C3b(*(39–45)(46)

One method used by the RPE to preserve the integrity of the monolayer during increased apoptosis and damage is to expand the borders of neighboring cells to maintain contact. Consequently, this increases the polymegatheism and pleomorphism displayed in aging and damaged RPE cells. A study by Rashid and Bhatia showed a region-specific deterioration in the RPE during normal aging [(4)]; in addition, during AMD, there is reduced regularity and an increased cell size in central RPE ((4). Alpha-catenin (CTNNA1), a mechanosensory protein that exists in the cell wall of RPE cells [(63)] and interacts with F-actin and cadherins of the actin cytoskeleton, will be released from adheren junctions into the cytoplasm of the cells and is a marker of cellular stress. A sign of early AMD has been shown to be the reorganization of intracellular auto fluorescent lipofuscin granules into aggregates, which were then released into the sub-RPE space ((47)). There is also an association with increased age and decreased phagocytic capacity of aged RPE cells and patients diagnosed with AMD. (48–50). By understanding the common convergent and divergent hallmarks separating normal and abnormal aging mechanisms, we can better understand signs of pathogenesis initiation earlier and preserve vision for longer.

Multiple rodent models have been used to study hallmarks of aging, since the rodent retina and RPE exhibit similarities to that of humans, even though mice lack a macula. While mice are not a perfect model to study ocular damage and stress associated with increased age(51), the study of the mouse visual system and the advent of genetic engineering tools have greatly benefitted our understanding of multiple disease processes(51–54). In this study, we used a natural aging model to study unique structural patterning in the RPE associated with age and RPE deterioration. Previous aging studies use relatively young animals and compare them with animals that are ∼1 to 2 years old; however, this excludes a crucial demographic that may better capture overall progression of ocular differences due to aging that recapitulates disease in humans. Additionally, many of these studies focus on retinal dysfunction and may miss critical changes in RPE structure that are also predictive of visual function outcomes. Therefore, in this study we used a C57BL/6J mouse model divided into three groups ranging from young (P60-180) to aged (P730+) to analyze the impact natural aging has on the RPE structure and function. Our comparative study is one of the few that includes this aged group and provides a more complete representation of aging RPE morphometric phenotypes that mirrors the aging process in humans. In this study, we sought to expand our previous work in describing the topological patterns of RPE dysmorphia with age by adding a correlative analysis of immune cell recruitment, stress-associated protein expression changes, and RPE dysfunction that accompany loss of retinal function over time(55). Further study of age-related, structural RPE heterogeneity and its molecular underpinnings are required to better target RPE-mediated initiation of age-related retinopathies.

## 2. Methods

### 2.1 Animals

Mouse housing, experiments, and handling were approved by the Emory University Institutional Animal Care and Use Committee. Studies were conducted in adherence with Association for Research in Vision and Ophthalmology (ARVO) and followed guidance and principles of the Association for Assessment and Accreditation of Laboratory Animal Care (AAALAC). C57BL/6J (WT) mice were maintained on a 12-h light/dark cycle at 22°C, and standard mouse chow (Lab Diet 5001; PMI Nutrition Inc., LLC, Brentwood, MO) and water were provided *ad libitum*. Animals were either purchased from Jackson Laboratories (JAX) directly or bred in-house for 3 generations or less from JAX breeding pairs. The mice used in each group were collected from different litters, and all samples displayed represent an independent animal; therefore, we expect no batch effects. The mice were managed and housed by Emory University Division of Animal Resources. Adult mice were euthanized using CO_2_ gas asphyxiation for 5 minutes followed by cervical dislocation. All mice used for this study were divided up into the following groups: Group 1 (Young: post-natal day 60-180); Group 2 (Middle-aged: post-natal day 365-729) ; Group 3 (Aged: post-natal day 730+).

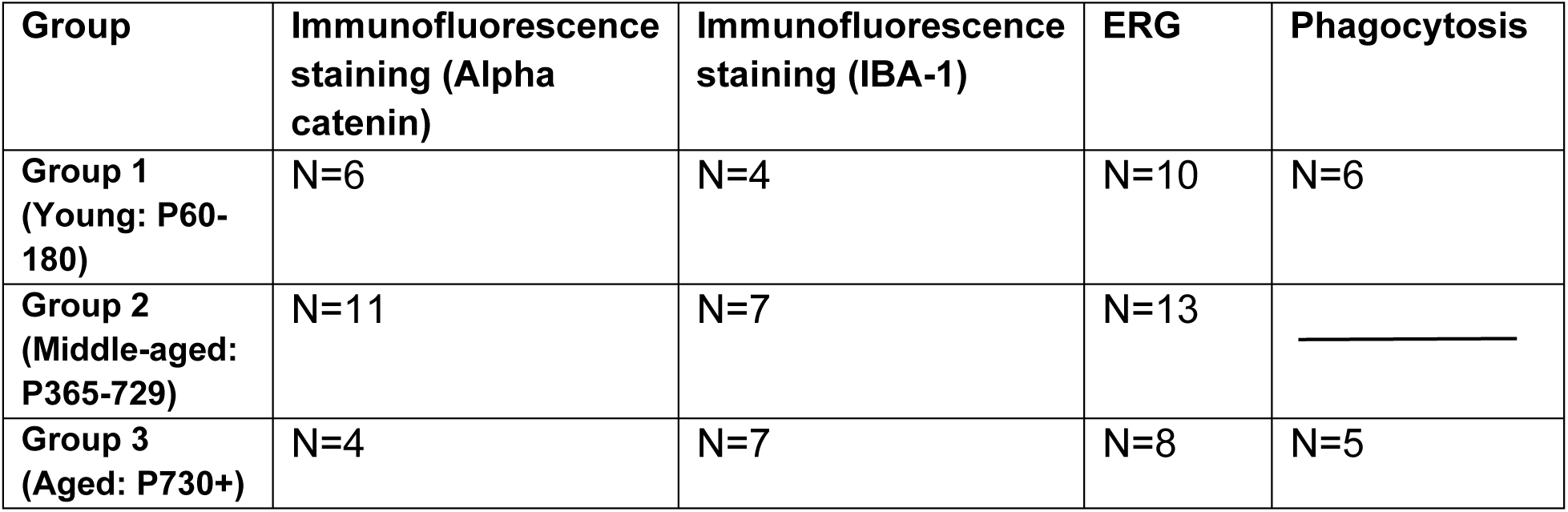

### 2.2 RPE and Visual Function Studies

#### 2.2.1. Electroretinograms (ERGs) – a-, b-, and c-Waves

Scotopic and photopic electroretinograms were performed on mice that were dark-adapted overnight. Each mouse was anesthetized using intraperitoneal (IP) injections of 100 mg/kg of ketamine and 15 mg/kg xylazine (ketamine; KetaVed from Patterson Veterinary, Greeley, CO; xylazine from Patterson Veterinary, Greeley, CO). Once anesthetized, the pupils were dilated with proparacaine (1%; Akorn Inc., Ann Arbor, MI) and tropicamide (1% Tropicamide Opthalmic Solution, USP; Akorn Inc., Ann Arbor, MI or 0.5% Tropicamide Opthalmic Solution, USP, Sandoz, Princeton, NJ) eye drops, which were administered topically. Mice were placed on a heating pad under red light and function was analyzed with Diagnosys Celeris System (Diagnosys, LLC, Lowell, MA). Full field ERGs were assessed at the following stimulus intensities (0.001, 0.005, 0.01, 1, and 10 cd s/m^2^). Scotopic a-, b-, and c-waves were collected. Afterwards, mice were injected with reversal agent (0.5 mg/mL atipamezole, injection volume 5 µL per gram mouse weight; Patterson Veterinary, Greeley, CO) and placed individually in cages on top of heated water pads to recover.

### 2.2. Phagocytosis

Murine eyes were enucleated and placed in glass tubes of “freeze-sub” solution of 97% methanol (Fisher Scientific A433p-4) and 3% acetic acid that were chilled with dry ice. Tubes were placed in -80°C for at least four days to dehydrate the tissue. After at least four days, tubes were allowed to reach room temperature before the eye samples were placed into tissue cassettes (Fisher Scientific, Catalog # 15200403D). The cassettes were then placed in 100% ethanol for 20 minutes, and then fresh 100% ethanol for another 20 minutes. Next, the cassettes were placed in xylene (Fisher Scientific X3S-4) for 20 minutes and then in fresh xylene for another 20 minutes. Afterwards, the cassettes were placed in a paraffin bath for 45 to 60 minutes before being transferred to a fresh paraffin bath for another 45-60 minutes. Eyes were then embedded in paraffin and sectioned for immunofluorescence.

The paraffin sections were then deparaffinized and rehydrated with xylene and decreasing concentrations of ethanol, then finally in Tris-buffered saline (TBS) (#1706435; Bio-Rad). The slides were covered in a blocking solution made up of in Tris-buffered saline (#1706435; Bio-Rad) with 0.1% (vol/vol) Tween 20 (pH 6.0) (BP337-100; Fisher Scientific) (TBST)] with 2% Bovine Serum Albumin (BSA) [catalog #BP9703-100].The primary antibody is then added to the blocking solution and put on the slides overnight at room temperature in a humidified chamber. The next day, the secondary antibody is added to the blocking solution. The slides were washed three times in TBST, then the secondary antibody is placed on the slides and incubated for four hours. Slides were washed and nuclei stained before mounting in flouromount G (catalog #0100-01; SouthernBiotech, Birmingham, AL, USA).

### 2.3. RPE Flat mount Preparation for ZO-1/Alpha Catenin Immunofluorescence

Murine eyes were enucleated and placed in zinc and formaldehyde (Z-fix) (Anatach, Battle Creek, MI Catalog # 622) for fixation for 10 minutes at room temperature. Afterwards, the eyes were washed five times in Hanks’ Balanced Salt Solution (HBSS Cat #14025092 Gibco by Life Technologies, Grant Island, NY) and stored at 4°C for up to 24 hours before dissection. RPE flat mounts were dissected and prepared as previously described in Zhang et al. (2021)(56). After removal of retina, each RPE flat mount was individually transferred into a well created by attaching a silicone gasket (Sigma Aldrich #GBL665104-25EA) to a SuperFrost Plus microscope glass slide (Fisher Scientific #12-550-15). Flat mounts were incubated in 300 µL of blocking buffer (3% (W/V) bovine serum albumin (BSA) (Catalog #BP9703-100) and 0.1% (V/V) Triton X-100 (Sigma) in HBSS (Fisher Scientific Catalog # MT21023CV) for 1 hour at room temperature in a humidified chamber. Primary antibodies (anti-ZO1 EMD Millipore Cat #MABT11 and anti-CTNNA1 (Alpha Catenin) Abcam Cat #ab51032) were diluted and pre-blocked in the blocking buffer for 1 hour prior to being applied to the flat mounts. Flat mounts were incubated in primary antibody overnight at room temperature. The primary antibodies were aspirated, and the flat mounts were washed five times with wash buffer (HBSS and 0.1% V/V Triton X-100). Secondary antibodies (see Table 1) were diluted and pre-blocked in blocking buffer for 1 hour at room temperature before being applied to the flat mounts overnight at room temperature. The flat mounts were then rinsed three times with the nuclei stain, Hoechst 33258 (Thermo-Fisher H3569, Waltham, MA), in the wash buffer, followed by two additional washes with the wash buffer. Afterwards, the wash buffer was aspirated, the gasket was removed from the glass slide, and the flat mounts were mounted with Fluoromount-G (SouthernBiotech; Catalog #0100-01; Birmingham, AL) and covered with a 22X40mm coverslip (Thermo Fisher #152250). Flat mounts were allowed to dry overnight on flat surfaces in the dark.

**Table 1:** Antibodies Used[1] [2].

| Antibody | Antibody type | Vendor + Catalog # | Concentration |
| --- | --- | --- | --- |
| Rat Anti-ZO1 | Primary | EMD Millipore, Catalog # MABT11 | [1:200] |
| Rabbit Anti-CTNNA1 (Alpha Catenin) | Primary | Abcam, Catalog #AB51032 | [1:500] |
| Rabbit Anti-Iba1(ionized calcium binding adaptor molecule) | Primary | Abcam, Catalog #ab178847 | [1:1000] |
| Mouse anti- Rhodopsin | Primary | Abcam, ab3267 | [1:250] |
| Rabbit anti-Best1 | Primary | Abcam, ab14927 | [1:250] |
| Pentahydrate (bis-Benzamide) Hoechst 33258 | Primary | Thermo-Fisher Catalog #: H3569 | [1:250] |
| Donkey anti-Rat (AF488) | Secondary | Life Technologies, Catalog # A21208 | [1:1000] |
| Donkey anti-rabbit (AF568) | Secondary | Life Technologies, Catalog # A10042 | [1:1000] |
| Donkey Anti-Mouse | secondary | Life Technologies Catalog #A21202 | [1:1000] |

### 2.4. RPE Flat mount Preparation for ZO1/Iba1 Immunohistochemistry

Enucleated murine eyes were fixed in 4% paraformaldehyde (PFA; 16% solution stored under argon from Electron Microscopy Sciences Catalog # 15710) diluted in 1X PBS (Fisher Scientific #50-980-487 and Corning 46-013-CM) for 1 hour at room temperature. Afterwards, the eyes were washed five times in Hanks’ Balanced Salt Solution (HBSS Cat #14025092 Gibco by Life Technologies, Grant Island, NY) and stored for up to 24 hours before dissection at 4°C. Following dissection, RPE flat mounts were individually transferred into a well created by attaching a silicone gasket (Sigma Aldrich #GBL665104-25EA) to a SuperFrost Plus microscope glass slide (Fisher Scientific #12-550-15). The flat mount was then rinsed with HBSS and incubated in blocking buffer made with HBSS with 5% bovine serum albumin (Catalog #BP9703-100) (W/V) and 3% Triton X-100 (Sigma) (V/V) for 2 hours at room temperature. Primary antibodies (anti-ZO1 EMD Millipore Cat #MABT11 and anti-Iba1 [EPR16589] (ab178847)) were diluted and pre-blocked in the blocking buffer for 1 hour prior to being applied to the flat mounts. After 2 hours, the blocking buffer was aspirated and the flat mount was incubated in the primary antibodies overnight. The next day, the primary antibody solution is aspirated, and the flat mount was rinsed five times for 2 minutes each with the wash buffer (HBSS and 0.3% Triton X-100 (V/V)). Secondary antibodies (see Table 1) were diluted and pre-blocked in blocking buffer for 1 hour at room temperature prior to flat mount application. Flatmounts were incubated in secondary antibodies overnight at room temperature. Each flat mount was rinsed three times with the nuclei stain, Hoechst 33258 (Thermo-Fisher H3569), in the blocking buffer and then two more times with the wash buffer. The gasket was removed from the glass slide and the flat mounts were mounted with Fluor mount-G (SouthernBiotech; Catalog #0100-01; Birmingham, AL) and covered with a 22X40mm coverslip (Thermo Fisher #152250).

### 2.5. Confocal Microscopy

The Nikon Ti2 with A1R confocal scanner microscope was used for imaging. Processing software used for imaging was NIS Elements 5.2. Imaging was done in resonance mode at 1024×1024 with 8x averaging and a Denoise.ai filter was applied to the images. Lasers were 405, 488, 560, 640 nm. Images were collected using a 20x objective and usually 25 images were photomerged together with using Adobe Photoshop CS6.

### 2.6. CellProfiler for Morphology Analysis

CellProfiler, a free, open-sourced cell image analysis software, is designed to analyze different images through customizable scripts, or “pipelines”. A pipeline was created specifically to analyze the morphology of the retinal pigment epithelium cells in murine eyes (used with Cellprofiler version 4.2.5), specifically the area, eccentricity, and radius. The pipeline first converted the staining of ZO-1 (green), Alpha Catenin (red), and Hoescht 33258 (blue) to gray, inverted the image, identified the primary objects in the image, collected metrics of each individual cells (including the number of neighbors and eccentricity), and saved all morphometric analysis information of each individual cell to a spreadsheet that can be exported from the program for analysis.

### 2.7. Imaris for Alpha-catenin Intensity

The intensity of cytoplasmic alpha-catenin was analyzed using Imaris software by Bitplane. Maximum intensity projection images of each RPE flat mount were processed using IMARIS 9.6 (Bitplane, Inc.) in which individual cells were segmented, identified, and quantified morphologically. Prior to converting and uploading images to Imaris, the corneal flaps and optic nerve heads were removed via the crop tool in Photoshop. Cropped flat mounts were uploaded to Imaris and segmentation was customized based on target cell characteristics. The Imaris software allowed for thresholding based on cell size and incorrectly segmented cells or artifacts were manually rejected. ZO-1 was used to segment each RPE cell and cells were filtered based on alpha catenin intensities in the cytosol.

### 2.8. Iba1 Quantification

Sub-retinal immune cells were manually counted using the Photoshop count function (Adobe Photoshop, Version 27.4.0 release) by three independent, masked, observers.

### 2.9. Statistical analyses

Statistical analysis was conducted using Prism 9.1.0 on Mac OS X Version 7 (GraphPad Software, Inc., La Jolla, CA, USA). All statistical tests used are summarized as mean +/- standard deviation (SD) and each individual statistical test is listed in the figure legends. A p-value <0.05 was considered statistically significant. Demographic distributions and sample sizes are summarized in Table 1. Each member of every group was an independent mouse.

## 3. Results

### Natural aging of the retinal pigment epithelium resulted in ectopic localization of structural protein, alpha catenin

To examine if increased age results in changes in alpha-catenin expression levels or distribution within RPE cells, we stained RPE flat mounts from young and aged mice. We found that with increased age, alpha catenin expression was more prevalent in the cytoplasm of RPE cells of aging mice compared to young mice [Figure 1]. The oldest mice showed a 1.9-fold increase in cytoplasmic alpha-catenin signal than the youngest mice; while middle-aged mice showed a 15% percent increase compared to the youngest group. This increased accumulation coincided with increased numbers of enlarged, multinucleated RPE cells (see white arrows, Figure 1). Enlarged cells were mainly concentrated in the central portion of the RPE sheet compared to the periphery. However, peripheral cells in the aged group also showed signs of cytoplasmic alpha-catenin. These changes in alpha-catenin localization were a sign of RPE stress as the cells undergo structural modification in response to age-related damage.

**Figure 1:**
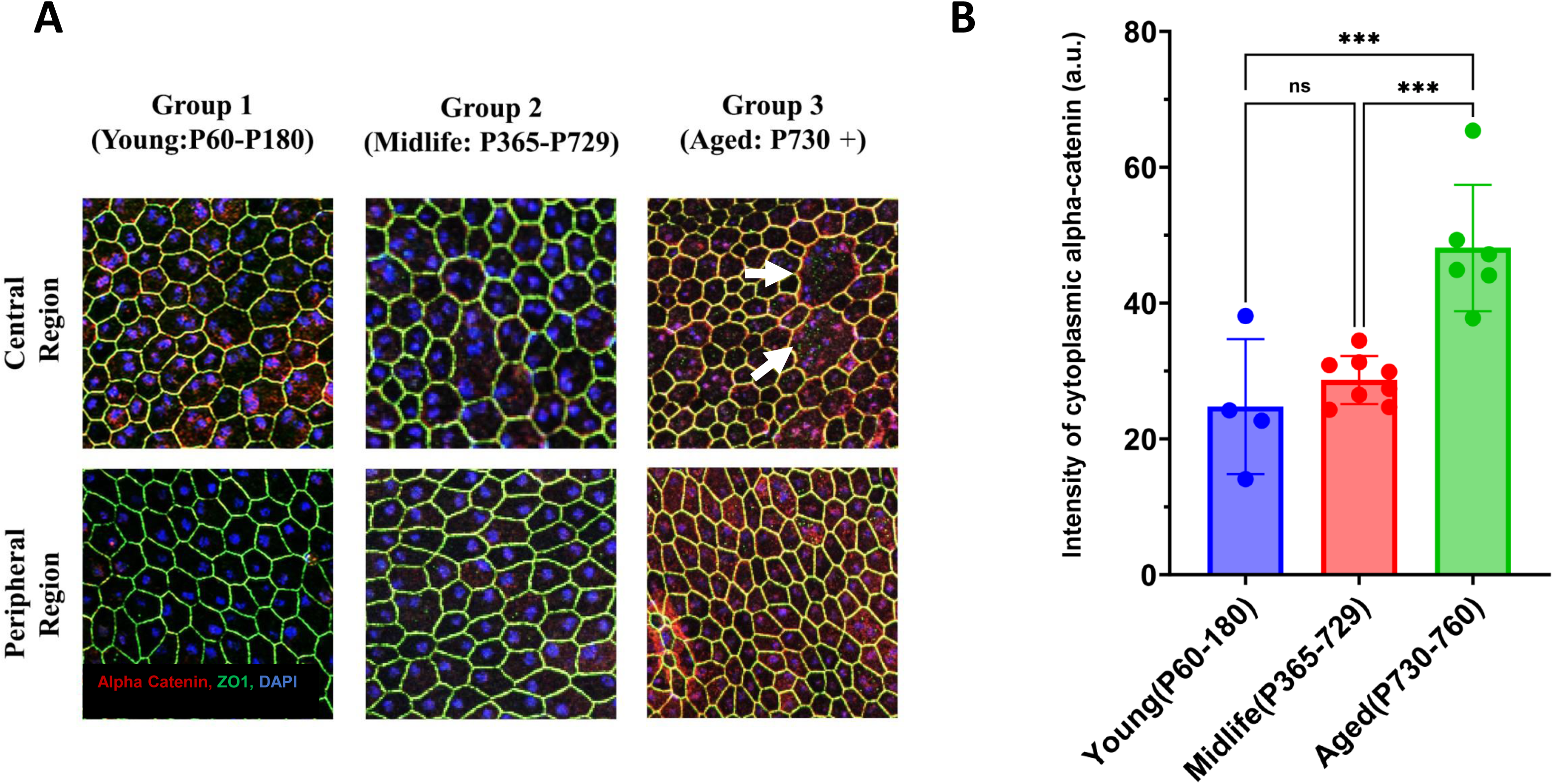
Natural aging of the retinal pigment epithelium resulted in ectopic localization of the structural protein, alpha-catenin. RPE flat mounts from animals in Young (P60-180), Middle-aged (P365-729), and Aged (P730+) were collected and stained for anti-alpha catenin [1:500; red], anti-ZO-1[1:200; green], and DAPI [blue]. The figure shows a representative image of cytoplasmic localization of alpha-catenin into the cytoplasm of cells that exhibit atypical morphology [Figure 1A]. White arrows show examples of enlarged RPE cells with cytoplasmic expression of alpha-catenin. The prevalence of these cells increased in the aged group and is quantified in Fig 1B. n=4-8 animals/group. One-way ANOVA was used for analysis. Error bars: SD * Represents p value <0.05; ** represents p value <0.01; *** represents p value <0.001; **** represents p value <0.0001. Sample size: 4-8 animals/group.

### Cytoplasmic Alpha-catenin localization has regional distribution patterns and is highly expressed centrally in aging animals

The morphometric analysis of regional subcategories when averaged across the entire flat mount, showed no statistically significant differences among age groups [data not shown]. We hypothesized that by comparing RPE morphology by regional subcategories there would be a significant difference in morphometric features between age groups. To test this, we made five circular, concentric, partitions around the optic nerve expanding into the periphery. When these regional subcategories were compared between age groups, an interesting trend was highlighted. When comparing regions that were proximal to the optic nerve, there was a statistically significant increase of alpha-catenin expression that stratified the groups by age [See figure 2B]. This trend was further evident when assessing eccentricity and area mean. However, there was no significant difference among groups when assessing the diameter of the cells using Cell Profiler. We found that cytoplasmic alpha-catenin was increased in the aged group with regions proximal to the optic nerved containing the highest expression of alpha catenin. Additionally, the overall area and eccentricity (the ratio of the distance between the foci of the ellipse best fit to the cell to its major axis length) of the cells proximal to the optic nerve were smaller in the aged group compared to the young group. These data align with previous studies that show central RPE cells exhibit more cell death and stress signals than peripheral RPE cells [(23)]. The loss of their neighbors in the central RPE sheet may be compensated for by increased eccentricity of peripheral RPE cells possibly to conserve central RPE cell density and maintain the properties of the retina-RPE interface [Figure 2B and 2C].

**Figure 2:**
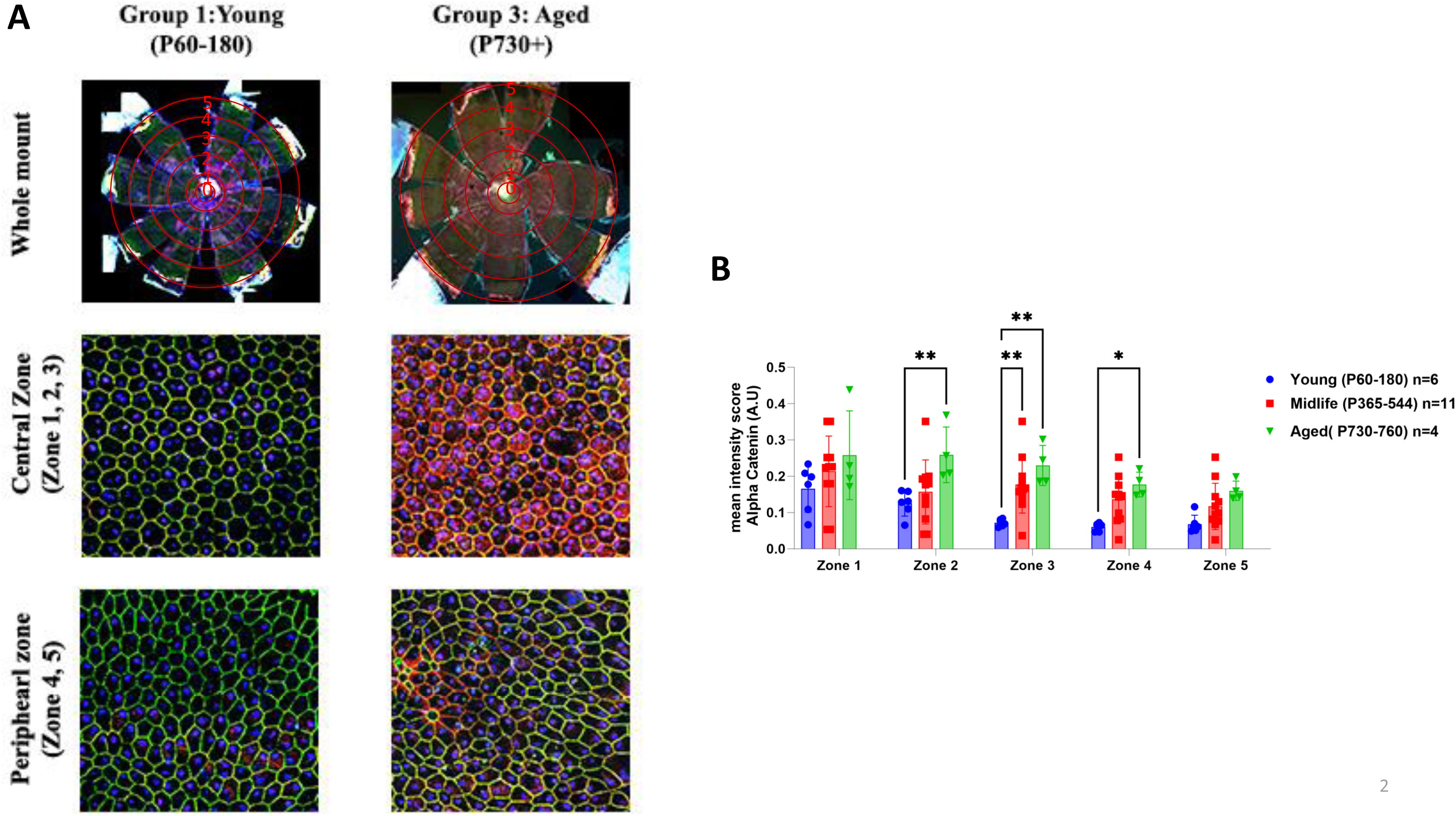
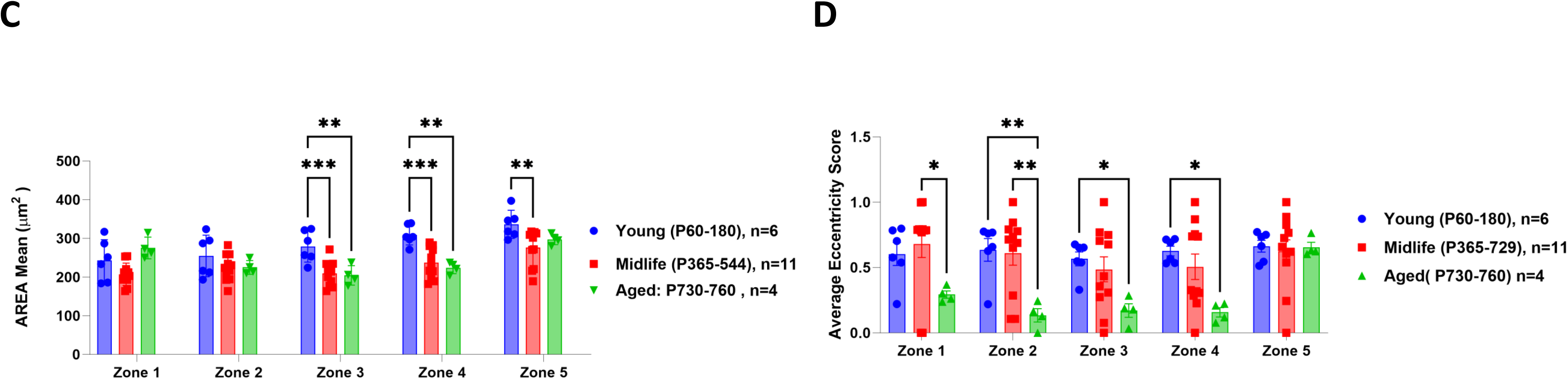
Cytoplasmic Alpha-catenin localization displays regional distribution patterns and was highly expressed centrally in aging animals. RPE flat mounts were segmented into concentric zones around the optic nerve and cropped for segmentation using CellProfiler. Zone locations are shown in representative image in Figure 2A. Multiple parameters were analyzed including the mean intensity of alpha catenin within the cytoplasm of the RPE cells (See figure 2B),, average area of cells(Figure 2C) and RPE cell shape (eccentricity) (Figure 2D). N=4-11/group. Analysis: Two-way ANOVA withTukey’s comparision (2B,2C and 2D); error bars: SD, *=p<0.05, **= p<0.01, ***= p<0.001.

### Inflammatory cell deposition within the RPE sheet increased with advanced age

To assess whether subretinal immune cell deposition was increased in natural aging of mice, we quantified the total number of IBA-1 positive cells in the RPE sheets of young and aged mice. We found that there was an overall 8-fold increase of IBA-1+ immune cells within the RPE sheets of mice in the aged group (aged) compared to younger groups [see figure 3A and B]. Additionally, the deposition of immune cells was 2-fold higher in the mid-periphery of the RPE within the aged group represented by zone 3 and 4 [see figure 3C] compared to the youngest group.

**Figure 3:**
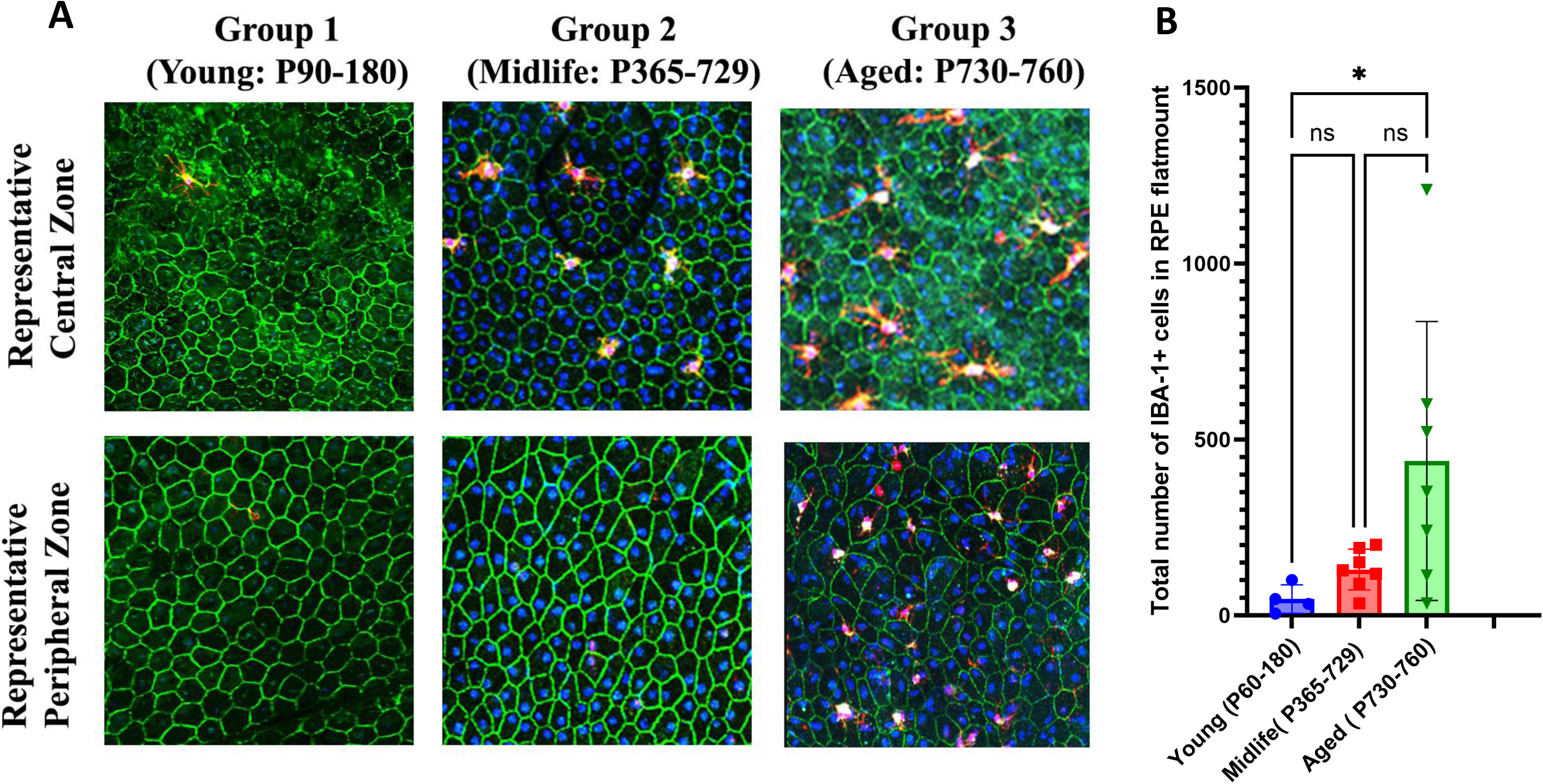
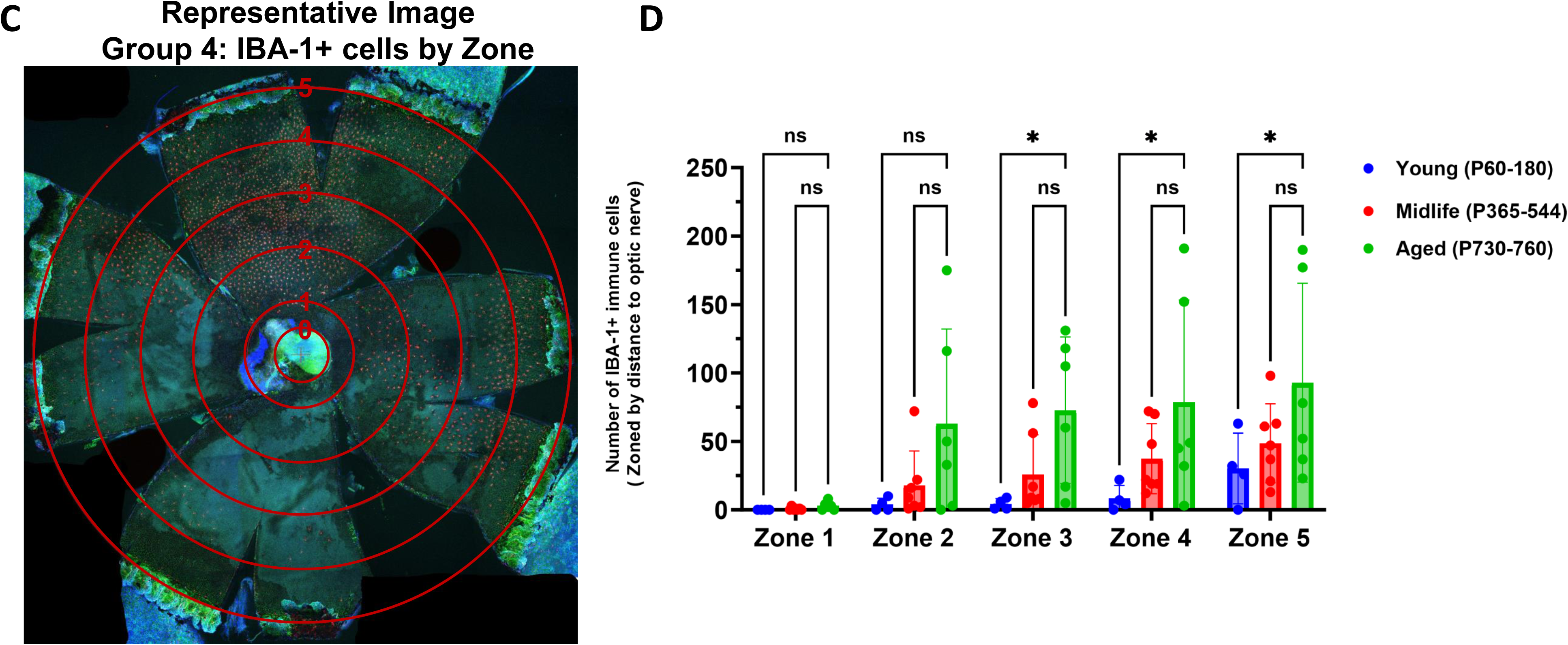
Inflammatory cell deposition within the RPE sheet increased with increasing age. RPE flat mounts were stained with an inflammatory cell marker, IBA-1[1:1000; red], ZO1(1:200; green], and Hoechst 33258 [blue]. Representative images show increased deposition of IBA-1 positive cells both centrally and peripherally in the aged group compared to the youngest in Figure 3A (group 1). The total IBA-1 positive cells were counted using Imaris [Figure 3B]. The flat mounts were then segmented into zones as previously mentioned and quantified with CellProfiler pipeline per zone. Quantification of these results are shown in Figure 3C and 3D. N=3-7 animals/group. Analysis: and Kruskal-Wallis with Dunn’s Correction test (3B) and Two-way ANOVA with Tukey’s comparison test(3D); error bars: SD *=p<0.05, **= p<0.01, ***= p<0.001

### Loss of retinal and RPE function occurs with natural aging

Comparative analysis of functional output between age groups found that there was an ∼50% loss of function in both scotopic a-, and b-wave function between the youngest and aged groups. A more modest but significant reduction in scotopic a- and b-waves were observed between the middle-aged group and the aged group and the young and middle-aged groups. (29% and 22% loss, respectively). When assessing RPE function via c-wave amplitude analysis, only the aged group showed a significant reduction in function compared to the youngest group (24% loss), while there was no difference between the youngest and middle-aged group.

### Aging animals show modest retinal and RPE morphological irregularity

Mice in the aged group showed marginal differences in retinal architecture compared to the youngest group. At ∼500 to 1000 microns from the optic nerve on the superior side, there is a much as a 34% increase in retinal thickness (young group: average: ∼ 6.2um ± 0.8 µm; Aged group: average 8.3um + 0.87 um. Two-way ANOVA with Dunnett’s comparison test) in the aged group compared to the youngest group. This difference in regional retinal thickness appeared near the optic nerve on the superior portion of retina. Additionally, the retinas of older mice showed abnormalities of the RPE layer with enlarged cells compared to young animals and loss of cell-cell contacts between the RPE cells indicating irregularities in the cell structures (See Figure 5A: see white arrows; Figure 5B). There were also changes in the morphology of the inner and outer segments of the photoreceptors in the aged group compared to the younger groups. These data also suggest that there were isolated, regional changes that were correlated with aged.

**Figure 4:**
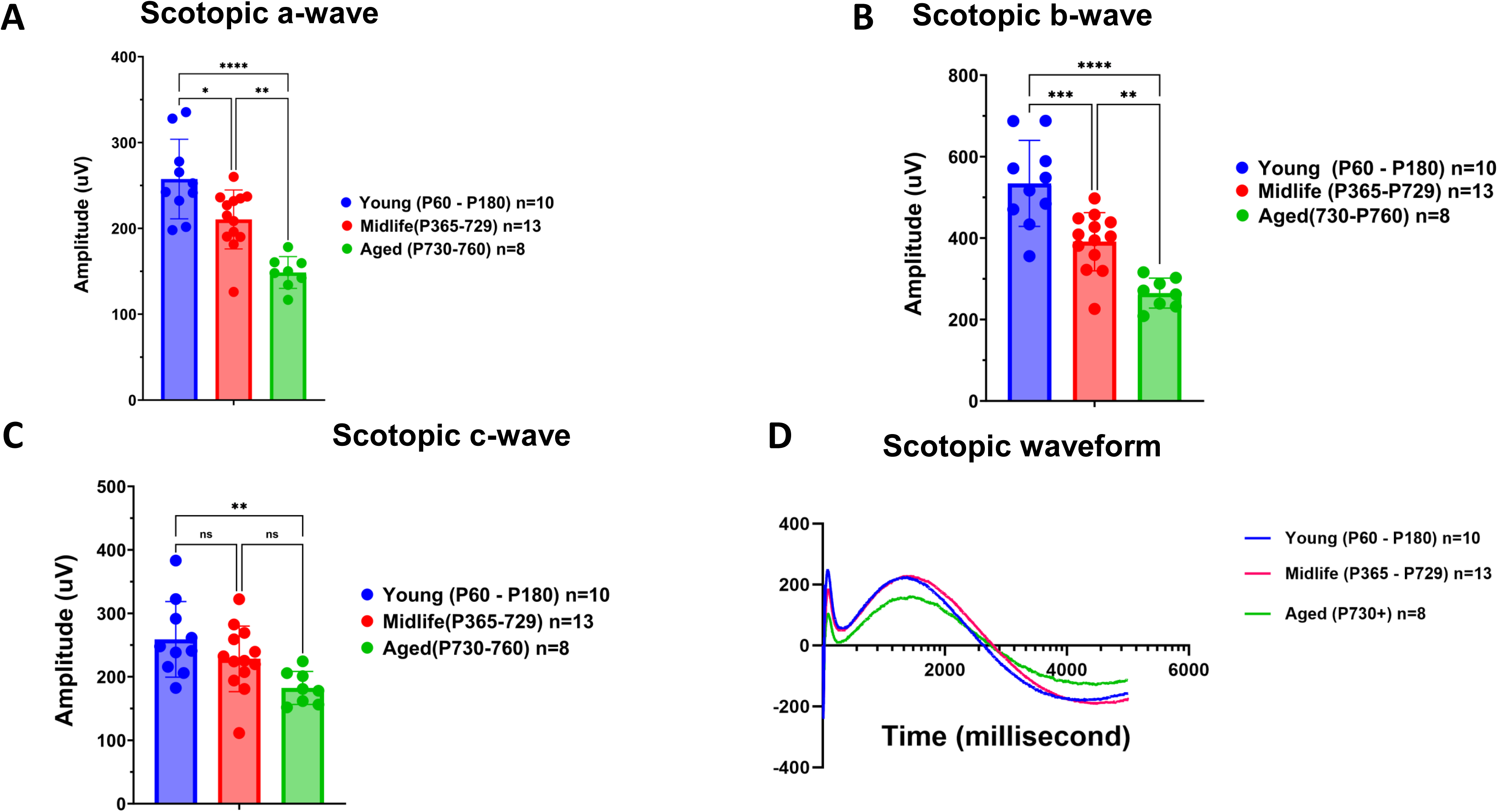
Significant loss of visual function with natural aging with moderate loss of function within the RPE. Raw electroretinogram waveforms from the young, middle-aged, and aged groups under scotopic conditions and after a 10-cd s/m^2^ light flash are shown (Panel D). At multiple flash intensities, there were significant differences between all groups and group 1 for scotopic *a-wave*, and scotopic *b-wave*, while the scotopic *c-wave* was only modestly significant between the young and aged groups. The significant reductions in ERG response, suggests that there were dysfunctional photoreceptors and bipolar cells. Additionally, the reduction in RPE c-wave response at 10Hz in aged group compared to young group suggests that the age-related changes in the RPE were affecting visual function, as well. One– Way ANOVA with Tukey’s comparison test*** represents p value <0.001; **** represents p value <0.0001 Samples sizes: 8-13 animals/group.

**Figure 5:**
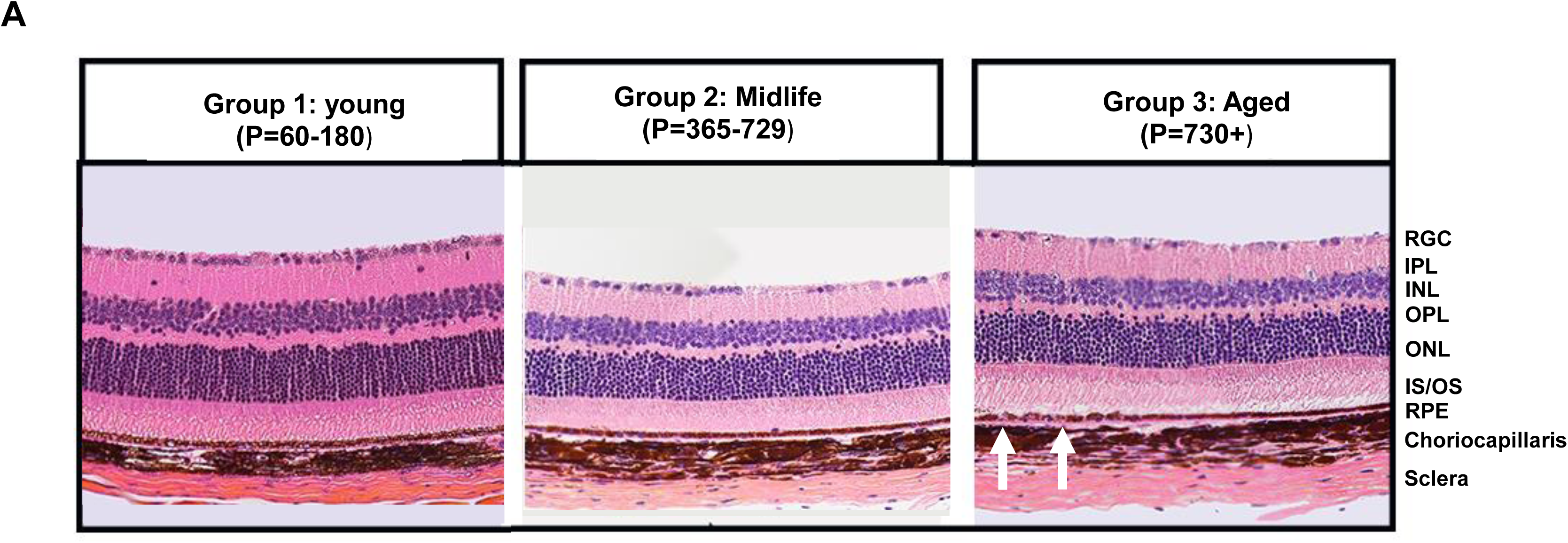
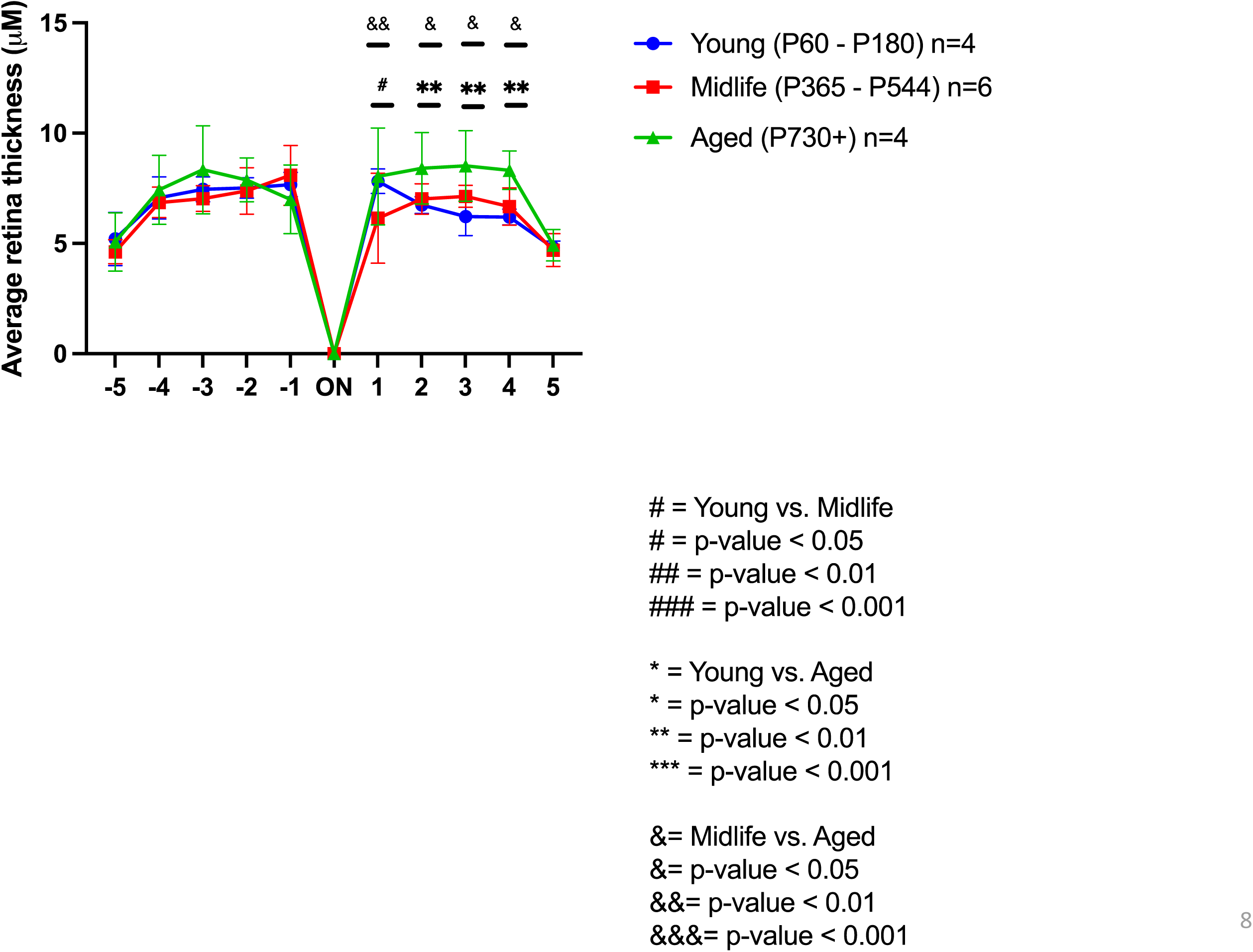
Aging animals showed modest signs of irregular morphology and retinal swelling in Hematoxylin & Eosin (H&E) staining compared to young animals. High magnification retina images (Figure 5A: panel 1-3) were shown for all 3 groups. The quantification of total retinal thickness is shown in figure 5B and shows regional changes in retinal thickness of aged mice compared to the youngest group. Two – Way ANOVA with Dunnett’s correction multiple comparisons test for retinal thickness analysis*, ^#^ = p value <0.05; **, ^##^ = p value <0.01; ***, ^###^ = p value < 0.001; ****, ^####^ = p value < 0.0001 (* symbols indicate significance between young group vs. aged group: # symbols indicate significant between the young and aged group. Samples sizes: 4-6 animals/ group)

### Natural aging resulted in retention of phagosomes within the RPE

A major function of the RPE is to provide cellular waste management of the photoreceptor outer segments via phagocytosis. Studies of RPE function in human donor eyes with AMD showed dysregulation of phagocytosis in the RPE eyes. Due to the significant loss of RPE function in the aged group compared to the young group, we evaluated RPE function via a rod outer segment phagocytosis assay. Based on previous studies, we hypothesized that there would be a reduction in phagosomes within the RPE of aging mice compared to the youngest (50) (57–59). However, we found an increase of ∼48% in phagosomes retained within the RPE of the aged group compared to the youngest group (young: ∼180 phagosomes/animal; aged group: ∼371 phagosomes/animal) (Figure 6A ad 6B).

**Figure 6:**
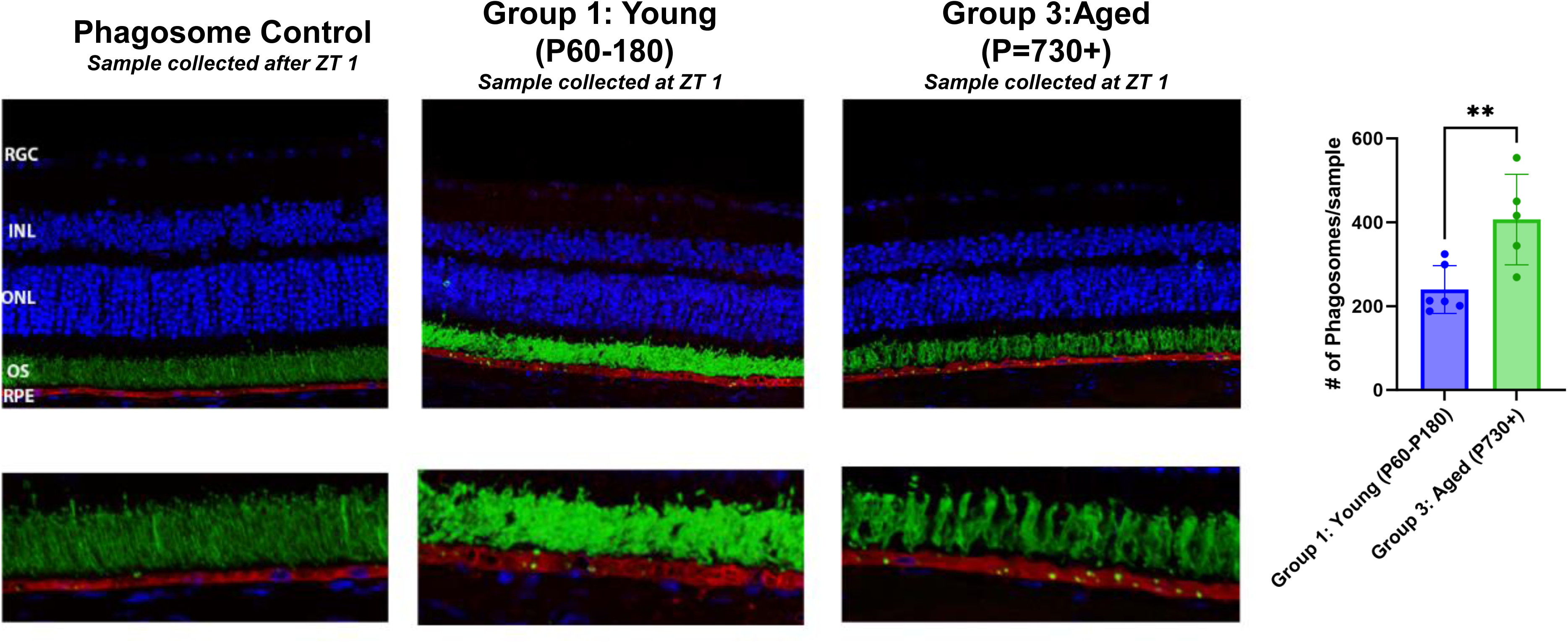
Natural aging resulted in retention of phagosomes within the RPE compared to young animals. Whole eyes were extracted from animals at Zeitgeber 1 [ZT1] within 1 hour of environmental light cue onset [ZT1] in each group. The sections were stained with Rhodopsin [green], Best1 [red], and DAPI [blue]. Representative images of a phagosome accumulation in the healthy control animal group [note: sample collected outside of ZT1] (far left panel), young group (middle panel) and aged group (far right panel) from a that was collected outside of the maximal phagosome production time (note: young and aged group samples were collected after ZT1 or 1 hour after lights on (7AM)). Rhodopsin containing phagosomes within the RPE were counted manually by three blinded observers and quantified. Results showed increased phagosomes within the RPE of the oldest group compared to the youngest animals or the ZT control suggesting that there may be aberrant turnover or maturation of the RPE phagosomes in aging mice. N=5-6/group. Analysis: Unpaired t-test ; error bars: Standard deviation *=p<0.05, **= p<0.01,***= p<0.001.

## 4. Discussion

The goal of this study was to define the changes in the RPE morphology during natural aging in the C57BL/6J mouse. In this article, we sought to systematically dissect these topological characteristics to look for discrete regional changes that may account for loss of central vision during natural aging. Previous work from our lab analyzed changes in RPE organization and topology in aging animals using morphometric analysis ((55)). In that study, we described RPE topology as relatively conserved characteristics between young and aged animals: 1) cells at the periphery radiated outward along an axis and show high levels of variability in cell shape, and 2) the cells located at the central to mid-peripheral RPE sheet are densely packed and show little change in density over time, possibly due to cells from the periphery elongating and pushing into the center. Thus, it is the accumulation of subtle RPE changes over time that are most correlated with the detrimental effects seen in aging. While our lab has explored multiple methodologies by which to optimize tracking subtle changes in RPE morphology that correlate with genetic and age-related manifestations of vision loss and pathology(60–62), our understanding of very early signs of RPE distress and damage remains elusive. This study builds on these methods, by concurrently evaluating increased cytosolic alpha catenin, increased pleomorphism and polymegathism in central RPE cells, reduced phagocytic capacity, and increased immune cell recruitment to the RPE as predictive of decreased overall RPE function with aging.

The RPE consists of a monolayer of post-mitotic cells that do not readily proliferate, consequently, the RPE has adopted other stress responses to accommodate for loss of cell density with age [(24)]. One of these methods is to increase the *en face* cell area to maintain cell-cell contact with healthy neighbors. Release of alpha-catenin (CTNNA1), a force-sensing protein that [(63)] interacts with F-actin and cadherins of the actin cytoskeleton, from adherens junctions has been reported to be associated with cellular stress in RPE cells. Multiple studies have validated the role of alpha-catenin in controlling actin skeleton dynamics via its ability to bind actin bundles and acts as a stabilizer at sites of adherens junction force-sensors [(63,67–70)]. Missense mutations in *CTNNA1*, which encodes alpha-catenin results in a distinctive “butterfly-shaped dystrophic “phenotype in RPE patterning leading to pathology. [(64)(12,23,65,66)]. We first evaluated the localization of alpha-catenin in conjunction with changes in the transmembrane tight junction protein, Zonula Occludins 1 (ZO1) in young, midlife, and aged mice as a metric for early signs of RPE damage repair. We found that in young mice most cells were of uniform cell size and shape and alpha catenin was restricted to the cell borders where it is likely bound to the actin cytoskeleton. When comparing general features of flat mounts by age group, animals in the aged group expressed higher levels of cytoplasmic alpha catenin via immunofluorescence than the young group [See Figure 1A-B]. This suggests that the aged group displays a higher degree of disassembly of adherens junction protein complexes releasing alpha-catenin into the cytoplasm where it may affect transcriptional regulation via beta-catenin binding. However, when assessing cell area and eccentricity (level of shape irregularity) when averaged over the entire flat mount, there were not significant differences between groups (data not shown).

Previous studies have examined RPE cell morphometrics in the aging C57BL/6J mouse and found that there were regional differences in RPE cell morphology [(23,55,71). Additionally, these studies highlight the degree of stress exerted at the cell borders contribute to regional heterogeneity in RPE structures with central and mid-peripheral RPE cells manifesting morphological and functional damage markers first [(16,77–81, 66)]. Considering the degree of heterogeneity that the RPE can display, we performed a regional analysis of cytoplasmic alpha catenin expression, cell area, and eccentricity to look for damage indicators. When stratifying by region, we found that zones 2, 3, and 4 were significantly enriched in cytoplasmic alpha catenin expression in the old group compared to the young [See figure 2A and 2B]. Additionally, in the aged group there was an increased degree of pleomorphoric cells (increased eccentricity) displaying polymegathism (non-uniform cell area) in zones 2, 3, and 4 in the old group compared to the young [See figure 2C-2D]. Del Priore et al, reported that apoptotic RPE cells increase with age and were prominently restricted to the central area near the fovea in humans [(23)] This may be due to the higher metabolic demands on central RPE, which must support a higher density of photoreceptors per cell. While mice lack a fovea, the photoreceptor dense region of the eye corresponds to zones 2, 3, and 4 radiating out from the optic nerve. These differences in metabolic demands may account for the variation in cytoplasmic alpha-catenin distribution seen in the aged group compared to the young regionally. This suggests a correlation between mislocalization of alpha-catenin to the cytoplasm and changes in RPE structure and size. An alternative explanation for the increased detection of cytoplasmic alpha catenin is the loss of melanosomes that occurs with increased age ((78). If melanosomes in the RPE are obstructing visualization of cytoplasmic alpha catenin, then this may explain why there appears to be more of it with aging as there is a loss of melanin with natural aging.

Next, we looked at recruitment of immune cells to the RPE sheet by staining RPE flat mounts for IBA-1+ cells. The increased subretinal infiltration of immune cells and dysregulation of inflammation associated genes has been linked to risk for AMD and damage in humans(79,80)(81,82). Genome-wide association studies and functional studies of the aging eye have shown that increased inflammation is a hallmark aging in the eye (33,88–92). Genetic models of AMD in mice suggest that dysregulation of microglia results in hallmarks that mimic AMD, i.e. RPE degeneration, atrophy of photoreceptors, and deposition of activated microglia in the subretinal space [(88)(89)]. It has been postulated that interactions between RPE cells and microglia/macrophages were involved in driving damage during AMD and other damage processes affecting the outer retina. Our study confirms this phenotype of increased deposition of subretinal immune cells and their attachment onto the apical face of the RPE overall in the oldest group(P730+) compared to the younger groups [See Figure 3AB]. Additionally, these cells show an increased presence within zone 3 and 4 in the oldest group, which corresponds with the mid-periphery to far periphery. Fewer immune cells deposited close to the optic nerve. The regional IBA-1+ cell deposition corresponds with the increased alpha-catenin expression and increased eccentricity data shown in Figure 2A-D, suggesting that there may be a correlation between RPE stress indicators, like cytoplasmic alpha-catenin, and increased regional accumulation of immune cells [See figure 3C-D]. As far as we know, we are the first to report age-related regional bias of subretinal microglia deposition patterns at the apical face of the RPE in mice. More study of subretinal immune cells during aging is needed to better understand the dynamics of their function and how expression of specific markers may elicit a pro- or anti-inflammatory program in the subretinal space [(83),(90)

Multiple groups have detailed how deterioration in visual function manifests with increased age in mice [(91–93)(94)]. This deterioration is hallmarked by reductions in visual acuity, functional cell responses using ERGs, and loss of visual performance and processing. When we assessed the visual function of young versus aged animals, we found that there is a significant decrease in function when compared to the young group (P90-P180) that begins relatively early on, appearing in midlife group (P365-P729) and continuing out to the aged group(P730+). There was a ∼50% loss of function in scotopic a- and b-waves between the young and aged group. When we assessed RPE by proxy of c-wave amplitude, there was only a significant loss of function between the young and the aged groups, which showed ∼30% decrease in function [See Figure 4A-C].

Taken together, these data suggests that there is a correlation between changes in early RPE stress indicators and significant loss of function between the young and old groups. Notably, the midlife group also begins to deviate significantly from the young group in both a- and b-wave function which corresponds with increased changes in region-specific polymegathism, polymorphism, and cytoplasmic alpha-catenin figure 2B-D. In the midlife group, the changes in visual function and RPE morphometrics precede increased subretinal immune cell infiltration or changes in c-wave in the midlife group.

We also examined retinal thinning and loss of cells over time. Interestingly, thickness measurements showed that there was an increase in retinal thickness in the aged group when compared to both the young and midlife groups. However, there was a regional component to this distinction, with the retina from the superior portion of the eye cup, within the mid-periphery, being thicker in the aged group compared to the young group [Figure 5B]. This may be indicative of edema or swelling of the retina proximal to the optic nerve due to cellular distress and may account for the regional increase in retinal thickness (95,96). There were also structural aberrations in RPE in the oldest group that do not appear in the younger three groups [Figure 4A denoted by white arrows]. Taken together, these data suggest that while the RPE is relatively resilient and manifest functional signs of damage later than the retina, the accumulation of structural changes in the RPE lead to loss of function and increased inflammation.

The accumulation of morphological changes paired with reduced clearance of oxidative stress by-products, lipid deposits and increased iron deposits can result in RPE dysfunction and are associated with natural aging [(97)(98–101). Finally, due to the loss of RPE functional electroretinogram output, we evaluated phagocytic activity of the RPE between the youngest and oldest groups. Phagocytosis is a major function of the RPE and is based on onset of light stimulus and circadian rhythm. We collected samples within 1 hour of light onset and stained them for rhodopsin to look at phagosome production between the youngest and oldest groups. We found that there were significantly more phagosomes in the oldest group than the youngest group (see Fig. 6). These data show that there is likely still dysfunction in the RPE, but we hypothesize the problem may be with phagosome turnover rather than a reduction in phagosome production or maturation. (59,102) It may be the incomplete proteolysis in the advance age RPE that contributes to the cellular stress and leads to loss of function and death of the RPE [(73). It would also partially account for the manifestation of regional differences, since the central RPE cells were responsible for recycling the photoreceptor outer segments of a higher density of photoreceptors. As a result, defects would be the most prominent and immediately detected in the central portion of the RPE before the periphery.

Limitations: This study is limited by the increased attrition rate of the study as animals age and drop out due to natural causes. Very few of the animals survived to be included in the oldest group, which resulted in an age and sex bias in the data. Additionally, the data for the RPE segmentation and immune cell counts was contingent on the quality of the fixation methods and minimization of artifacts during dissection. Another limitation of this study is that the morphometric analysis of RPE cells and immune cells is contingent on the preparation conditions used. Thus, only the cells that were adherent to the apical RPE surface at time of collection were assessed, which doesn’t account for non-adherent cells that interact with the RPE and may be involved in the damage response. Analysis of the flat mounts was restricted to the apical face, which may miss phenomena that contribute to pathology that lies on the basal face of the cells. Data analysis was also limited by the nature of the study only incorporating static, post-mortem analysis of cellular changes. If longitudinal, live-imaging data were incorporated, this would have complemented this study and given a more robust view of dynamic changes in the eye with natural aging. Additionally, studies of the dynamics of RPE cells and subretinal immune cells may differ between humans and mice during aging, thus incorporating human samples would have validated the assertions made in this study. We also used an inbred, C57BL6J mouse strain, as a result, the observations may differ in other inbred strains or in outbred mice. In this study, we did not delineate between microglia and macrophages which may play different roles in the onset of pathology associated with aging.

Future directions: This study highlighted the importance of spatiotemporal dynamics in both RPE cells and subretinal immune cells. By studying how subtle, accumulation of stress response protein expression (like that of alpha-catenin) correlate with changes subretinal immune cell deposition and changes in ocular function, we can better map the susceptibility to and progression of damage due to aging or disease. In future studies, being able to observe these subretinal immune and RPE dynamics in a time course using live imaging would be important to study and understand. Additionally, with the advent of spatial transcriptomics, we can take a deeper dive into cell responses in a more targeted manner and potentially identify candidates for intervention before visual loss occurs.

### Conclusion

In this study, we sought to further understand how regional changes in the RPE sheet due to aging may affect the functional and structural outcomes of the subretinal space. Our study found that RPE cells within the mid-periphery exhibit more eccentricity changes[Figure 2D], increased cytoplasmic alpha-catenin accumulation [Figure 1A-B, 2A-B], and increased deposition of IBA-1+ immune cells than the far periphery or central RPE cells in the advance age group [Figure 3A-D]. Our study also highlights how changes in regional stress responses and cell structure precedes notable loss of RPE function over time. These changes correlate with the loss of visual function via ERGs and defects in phagosome turnover in the RPE in the oldest group and were associated with pathology onset. Further study of potential mechanisms of RPE-immune cell communication during aging may give greater insight into targets for early intervention to preserve the sight of an increasing number of geriatric patients.

## Acknowledgments

Supported by Grants from the National Institutes of Health (R01EY028450, R01EY021592, P30EY006360, U01CA242936, R01EY028859, T32EY07092, T32GM008490); by the Abraham J. and Phyllis Katz Foundation; by grants from the U.S. Department of Veterans Affairs and Atlanta Veterans Administration Center for Excellence in Vision and Neurocognitive Rehabilitation (RR&D I01RX002806, I21RX001924; VA RR&D C9246C); and an unrestricted grant to the Department of Ophthalmology at Emory University from Research to Prevent Blindness, Inc.

## Notes

Funding: Supported by National Institutes of Health (NIH) grants R01EY028450, R01EY021592, P30EY006360, R01EY028859, T32EY07092, and T32GM008490, the Abraham and Phyllis Katz Foundation, VA RR&D I01RX002806 and I21RX001924, VA RR&D C9246C (Atlanta Veterans Administration Center for Excellence in Vision and Neurocognitive Rehabilitation), and a challenge grant to the Department of Ophthalmology at Emory University from Research to Prevent Blindness, Inc.

### Competing Interest Statement

The authors have declared no competing interest.

